# Reducing the CAV1-dependent trafficking of G6PC1 in the liver protects against the development of type 2 diabetes

**DOI:** 10.1101/2025.04.26.650768

**Authors:** Lorine Da Costa, Cécile Saint-Béat, Clara Bron, Arnaud Lang, Adeline Duchampt, Manon Micoud, Félicie Evrard, Gilles Mithieux, Amandine Gautier-Stein

**Author notes:** Corresponding author **Address for correspondence** Amandine Gautier-Stein, Université de Lyon, 7-11 rue Paradin, F-69008 Lyon, France.

## Abstract

Targeting hepatic gluconeogenesis is an efficient strategy to counteract the development of type 2 diabetes. Hepatic glucose production into the bloodstream is controlled by the GLUT2 transporter and a vesicular pathway dependent on Caveolin-1 (CAV1) involving G6PC1 location at the plasma membrane. We hypothesized that decreasing hepatic gluconeogenesis by targeting CAV1 specifically in the liver (L.*Cav1*^-/-^ mice) improves energy and glucose metabolism in diabetic mice. Here we show that the absence of hepatic CAV1 increases insulin sensitivity, glucose tolerance and decreases fasting hyperglycemia and hyperinsulinemia in mice feeding a high fat high sucrose diet (HFHS diet). The decrease in insulin sensitivity takes place also in L.*Cav1*^-/-^ mice feeding a standard diet (STD diet). Moreover, we demonstrated an improvement of glycemic control when hepatic *Cav1* is deleted in previously prediabetic mice. In parallel, the absence of CAV1 in the liver reduces body weight gain and lipid intestinal absorption. Together these findings highlight that the vesicular pathway of glucose production represents a promising therapeutic target in the prevention of type 2 diabetes.

## INTRODUCTION

The deregulation of endogenous glucose production (EGP) is a crucial point in the development of type 2 diabetes. Diabetic patients develop fasting hyperglycemia mainly due to excessive EGP. Particularly, hepatic glucose production (HGP) is higher in the post absorptive state and during fasting^1^. This mainly results from excessive gluconeogenesis^1–3^, which is inadequately suppressed by insulin in diabetic patients. The first-line anti-diabetic treatment prescribed is based on the suppression of hepatic gluconeogenesis via, in part, the reduction of *G6pc1* expression^4^. The *G6pc1* gene encodes for the catalytic subunit (G6PC1) of the glucose-6-phosphatase (G6Pase) complex. This complex includes the glucose-6-phosphate translocase (G6PT), which transports glucose-6-phosphate (G6P) from the cytoplasm to the endoplasmic reticulum lumen while G6PC1 catalyzes the last biochemical reaction of both glycogenolysis and gluconeogenesis by hydrolyzing G6P in glucose and inorganic phosphate^5^. The expression of G6PC1 is increased in the liver of diabetic patients^6^ and conversely, the absence of *G6pc1* in the liver of mice protects against the development of diabetes^7^. These studies demonstrate that G6PC1 plays a key role in the development of this disease and suggest that decreasing G6PC1 is a powerful strategy against type 2 diabetes development.

Hepatic glucose is produced in the endoplasmic reticulum (ER) and exported to the bloodstream by the facilitating transporter GLUT2 and a second pathway depending on vesicle trafficking ^8–10^. This pathway is able to compensate the lack of hepatic GLUT2 in the fasted state or upon glucagon stimulation ^11^. We recently demonstrated that this vesicular pathway involved in HGP implied the trafficking of G6PC1 to the plasma membrane, through a mechanism depending on Caveolin-1 (CAV1), the main protein forming caveolae. Indeed, the lack of CAV1 prevents G6PC1 transport from the ER to the plasma membrane and reduced glucose production from hepatocytes in the fasting state, independently of changes in G6PC1 levels ^8^.

To investigate the pathophysiological role of this new mechanism of glucose production, we developed a mouse model carrying an inducible liver-specific deletion of the Cav1 gene (L.Cav1^-/-^). We here assessed whether the deletion of hepatic Cav1 induced in adult mice might protect against the development of diabetes.

## MATERIAL AND METHODS

### Animals and diets

We used adult male L.*Cav1*^-/-^, L.*Glut2*^-/-12^ and wild-type (WT) mice. Sperm of homozygous mice carrying the mutant *Cav1* allele *(Cav1tm1c(KOMP)Mbp)* was obtained from the trans-NIH Knock-Out Mouse Project (KOMP) (https://www.komp.org/). The *Cav1tm1c* allele contains 2 loxP sites flanking the *Cav1* exon 2^13^. In vitro fertilization of C57BL6/J oocytes was performed by the animal facility of Lyon 1 University (Animaleries Lyon Est Specific Pathogen Free) to obtain B6.*Cav1*^ex2lox/lox^ mice that were further backcrossed with wild-type C57Bl6/J (Charles River Laboratories, France). To delete *Cav1* specifically in the liver, B6.*Cav1*^ex2lox/lox^ mice were then crossbred with transgenic mice expressing the inducible CRE^ERT2^ recombinase under the control of the serum albumin promoter (B6.SA^creERT2^/w)^14^.

Male B6.*Cav1*^*ex2lox/lox*^.SA^creERT2^ and B6.*Glut2*^ex2lox/lox^.SAcre^ERT2^ mice were injected intraperitoneally once daily (500µg/day during 5 days) at 8 weeks of age to induce *Cav1* or *Glut2* excision specifically in the liver and produced L.*Cav1*^-/-^ and L.*Glut2*^-/-^ respectively. Male wild-type C57Bl6/J (Charles River Laboratories, France) received the same tamoxifen injection at 8 weeks of age. All mice were housed on a 12-hour light/dark cycle in the animal facility of Lyon 1 University (Animaleries Lyon Est Conventionnelle and Specific Pathogen Free) in controlled temperature (22°C) conditions. Mice had ad libitum access to water and to a standard chow diet (STD, A04; SAFE; Augy, France) for 5 weeks (8 to 13 weeks of age). The diet was then prolonged or switched to a high-fat/high-sucrose diet (HFHS diet) (consisting of 36.1% fat, 35% carbohydrates (50% maltodextrine + 50% sucrose), 19.8% proteins)^15^ during 14 weeks (14 to 27 weeks of age). To delete *Cav1* in overweight and prediabetic mice, male B6.*Cav1*^ex2lox/lox^.SA^creERT2^ and WT mice were fed HFHS diet for 12 weeks and then received intraperitoneal tamoxifen injections (500µg/day during 5 days). These L.*Cav1*^-/-OB^ and WT mice were further fed a HFHS diet for 8 weeks. All the procedures were performed in accordance with the principles and guidelines established by the European Convention for the Protection of Laboratory Animals. All conditions and experiments were approved by the regional animal care committee (CEEA-55, Lyon 1 University, France, APAFIS #21202 and #28432) and the French Ministry of National Education, Higher Education and Research.

### Body weight and food intake

Body weight and food intake were measured once and twice per week, respectively, since weaning to the beginning of tolerance tests. As mice were housed in groups, total food intake was divided by the number of mice per cage.

### Euglycemic hyperinsulinemic clamp

Euglycemic hyperinsulinemic clamp were performed on 21-week-old mice fed a STD diet (13 weeks after tamoxifen injection). An indwelling catheter for insulin and glucose infusion was placed into the jugular vein under isoflurane anesthesia. Mice were allowed to recover for 8 days. After a 5h-fast, a 120 minutes euglycemic hyperinsulinemic clamp was conducted in awake, freely moving mice. A bolus of 1.25mU insulin was infused at the beginning of the clamp. Then, 3mU/kg/min insulin was infused during 120min. At the same time, 15% glucose solution was perfused at variable rates and adjusted every 10 minutes to keep stable glycemia. Glycemia was measured with glucometers from tail blood to adjust glucose flow rate.

### Lipids intestinal absorption

In 6-hour fasting mice, blood was withdrawn from the tail and collected in ethylene diamine tetraacetic acid (EDTA). Then Tyloxapol, a plasma lipase inhibitor, was injected in the caudal vein (500 mg/kg body weight diluted in saline solution). Mice were force-fed with 200µL of sunflower oil 15 minutes after tyloxapol injection. Blood was collected 0, 1, 2 and 4 hours after Tyloxapol injection from the tail. Blood samples were centrifuged 15min at 3000g, plasma were collected and stored at -20°C. Triglycerides were measured with a colorimetric kit in each plasma sample.

### Hepatocytes glucose production

Primary hepatocytes were obtained from 13-week-old WT and L.*Cav1*^-/-^ male mice after 16 h fasting by collagenase perfusion of the liver^16^. Hepatocytes were seeded into 6-well plates at 500,000 cells/well and cultured for 1 h at 37 C in a 5% CO2 air atmosphere in DMEM without glucose supplemented either with 10 mM lactate and 1 mM pyruvate. After centrifugation (5min, 280g), glucose was measured in the supernatant (extracellular glucose) and cell pellet (intracellular glucose) by enzymatic assay. The amount of glucose was normalized to the number of cells and expressed in pmol/million of cells.

### Glucose, Insulin and pyruvate tolerance tests

Glucose tolerance tests (GTT) and insulin tolerance tests (ITT) were performed in 6h-fasted mice at 25/26 weeks of age (after 12/13 weeks on HFHS diet)^17^. Pyruvate tolerance test (PTT) was performed in 16h-fasted mice at 20 weeks of age^18^.

### Body composition, energy homeostasis

After 14 weeks on HFHS diet, fat and lean mass were determined with nuclear magnetic resonance spectroscopy (Minispec LF90 II; BRUKER Society; ANR-11-EQPX-0035 PHENOCAN). For indirect calorimetry, oxygen consumption and carbon dioxide production were determined using a single chamber system and used for the calculation of energy expenditure (Phenomaster, TSE Systems GmbH). All procedures were performed on individually housed mice by the ANIPHY platform, SFR Lyon-Est from Lyon 1 University. Mice were placed in individual chambers 2 days before starting measurements, which were recorded during 2 consecutive days.

### Tissue sampling and biochemical assays

Tissue samples were collected from 30-week-old mice. After 6h or 16h of fasting, blood was withdrawn from the submandibular vein, collected in EDTA and conserved at -20°C. Then, mice were killed by dislocation, and tissues collected, frozen in liquid nitrogen, and stored at −80°C until needed for molecular and biochemical analyses. Plasma triglyceride and cholesterol concentrations were determined with colorimetric kits (Table 1). Plasma insulin concentration was quantified using Mouse ELISA kits (Table 1). Liver lysate was prepared at 4°C and immediately processed. Hepatic G6Pase activity was assayed at maximal velocity (20 mmol/l of G6P) at 30°C by complexometry of inorganic phosphate (Pi) produced from G6P. The phosphohydrolyzing activity toward β-glycerophosphate (20 mmol/l) was determined and subtracted from the total G6Pase activity in all cases to clear the specific G6Pase activity from the contribution of nonspecific phosphatases^19^. Glucose-6-phosphate and glycogen concentration was determined from liver lysate after deproteinization with perchloric acid (6% V/V) and neutralization with K_2_CO_3_ ^20^. The NADPH produced from the hydrolysis of G6P by G6PDH was then used to calculate G6P content ^20^. After neutralization of the lysate, glycogen was digested into glucose with *α*-amyloglucosidase, then glucose was measured as described ^21^. Total hepatic lipids were extracted by the method of Bligh and Dyer ^22^ and triglyceride and cholesterol contents were measured using a Biomerieux colorimetric kits (Table 1).

**Table 1.**
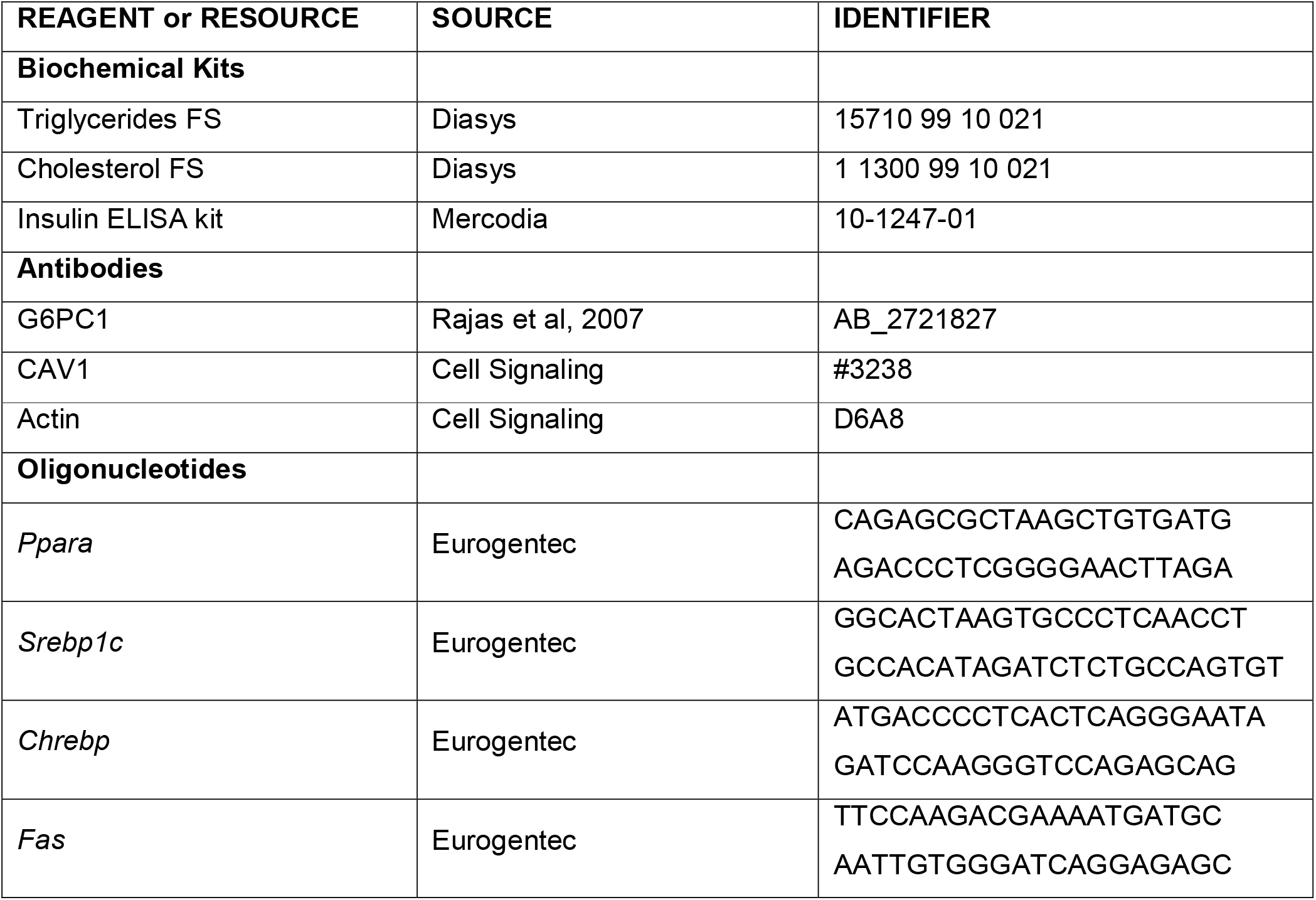
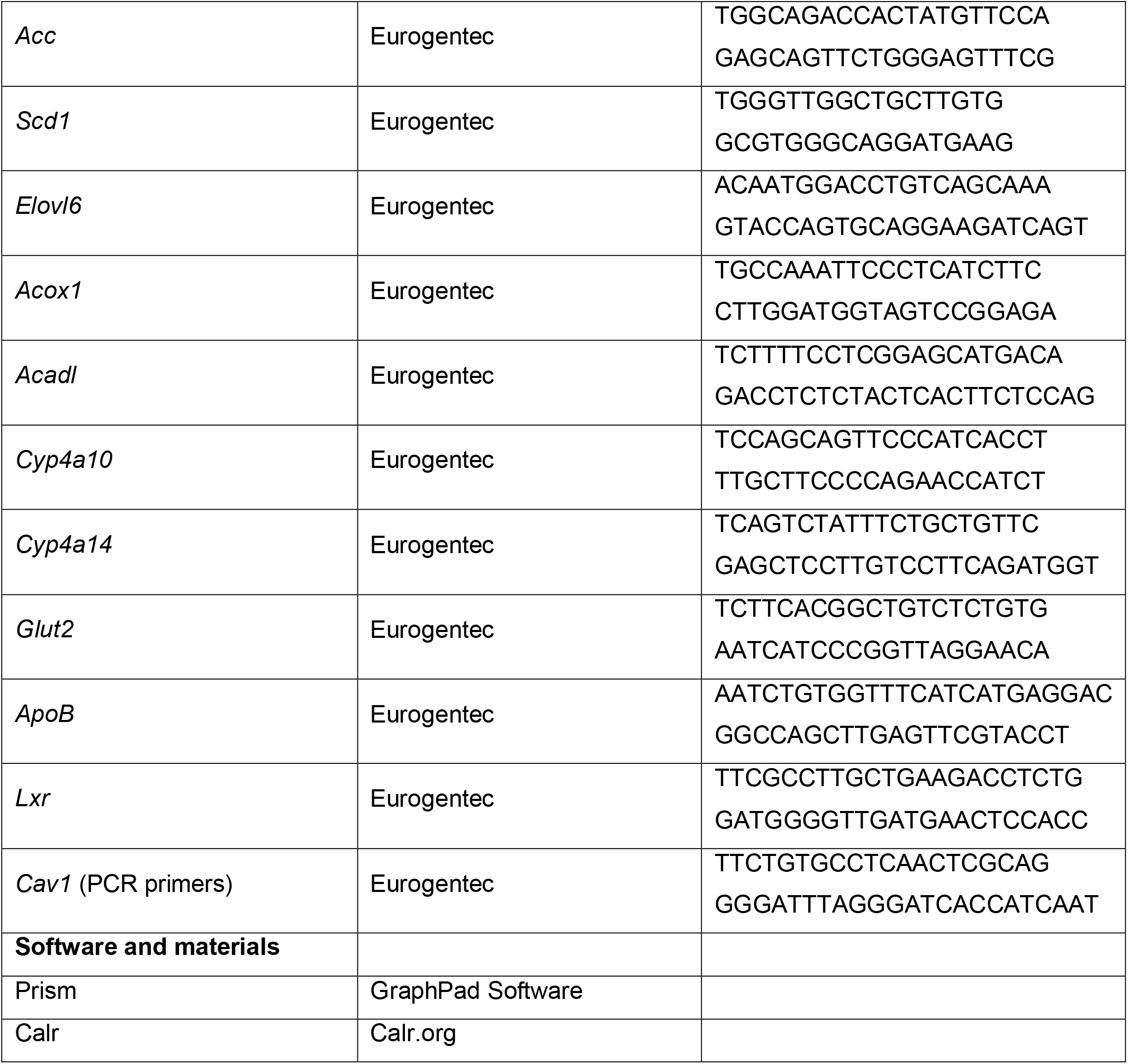

### Western-blot

Plasma membrane protein lysates were extracted from WT and L.*Cav1*^-/-^ liver as described previously^8^ and western blots were performed from 5µl of lysates using antibodies against G6PC1 (1:2000 ; Table 1). The relative amount of total proteins was used for normalization. Total protein lysates were extracted from WT and L.*Cav1*^-/-^ hepatocytes^8^. Western blots were performed from 30µg of proteins using antibodies against Caveolin-1 (1:1000; Table 1) and β-actin (1:5000; Table 1), which was used as loading control. Chemiluminescent signals were acquired with the Chemidoc Imaging System (Biorad) and quantified using Image Lab software.

### Gene expression analysis

A piece of 50 mg of frozen liver was homogenized with the Fast Prep® system and total RNAs were isolated according to the Trizol protocol (Invitrogen Life Technologies). Reverse transcription was performed on 1 μg of mRNA using the Qiagen QuantiTect Reverse Transcription kit. Real-time qPCRs were performed using sequence-specific primers with Takyon™ No Rox SYBR® Master Mix dTT Blue (Eurogentec). The expression of mRNA was normalized to the mouse ribosomal protein mL32 transcript (Rpl32) expression using the 2-ΔΔCt method. Primer sequences are reported in Table 1.

### Statistical analysis

Results are reported as mean ± standard error of the mean (SEM). Normal distribution was verified for all analyses. The effect of two independent variables on a dependent variable was analyzed using two-way ANOVA, followed by multiple comparisons, and corrected for multiple comparisons with a SIDAK test. Values were considered significant at *P < 0.05; **P < 0.01; ***P<0.001. Statistical analyses were performed with GraphPad Prism software. CalR was used for Figure 3G^23^.

## RESULTS

### The absence of hepatic CAV1 decreases hepatic glucose production

First, we checked that the deletion of *Cav1* was specific to the liver in L.*Cav1*^-/-^ mice. As shown in **Figure 1A**, PCR analysis revealed the presence of the recombined Cav1 allele in the liver but not in the intestine, kidney, white adipose tissue, lung and skin of L.*Cav1*^-/-^ mice. Hepatocyte CAV1 protein levels decreased by more than 80% in L.*Cav1*^-/-^ mice compared to WT mice 5 months after tamoxifen injection (**Figure 1B**). We previously reported that total *Cav1* deletion in *Cav1*^-/-^ mice prevented G6PC1 transport from the ER to the plasma membrane leading to a reduced HGP without affecting G6Pase activity^8^. Here, we showed a 40% decrease of G6PC1 levels in the plasma membrane of liver cells from 16h-fasted L.*Cav1*^-/-^ mice compared to WT mice with no change in G6Pase activity (**Figure 1C and 1D**). In line with previous results obtained in total *Cav1*^-/-^ mice, hepatocytes of 16h-fasted L.*Cav1*^-/-^ mice produced 30% less intracellular and extracellular glucose from pyruvate and lactate than WT hepatocytes, suggesting a decrease in *de novo* glucose production (**Figure 1E**). Of note, there was no difference in *Glut2* mRNA levels between the two groups (data not shown) and deleting *Glut2* in the liver of L.*Cav1*^-/-^ mice led to a further decrease in glucose production (L.*Glut2*^-/-^.L.*Cav1*^-/-^ hepatocytes produced 303443 ± 76574 pmol glucose/millions of cells). We confirmed the decrease in HGP *in vivo* by a pyruvate tolerance test. Glycemia was significantly lower 20 and 40 minutes after the intraperitoneal injection of pyruvate in 16h-fasted L.*Cav1*^-/-^ mice compared to WT mice, highlighting a decrease in HGP (**Figure 1F**). In parallel, glucose, G6P and glycogen levels were similar in the liver of 16h-fasted L.*Cav1*^-/-^ and WT mice fed a STD diet (**Figure 1G**). Taken together, these data show that the absence of CAV1 specifically in the liver reduces hepatic gluconeogenesis, in parallel to a reduction of G6PC1 levels at the plasma membrane.

**Figure 1:**
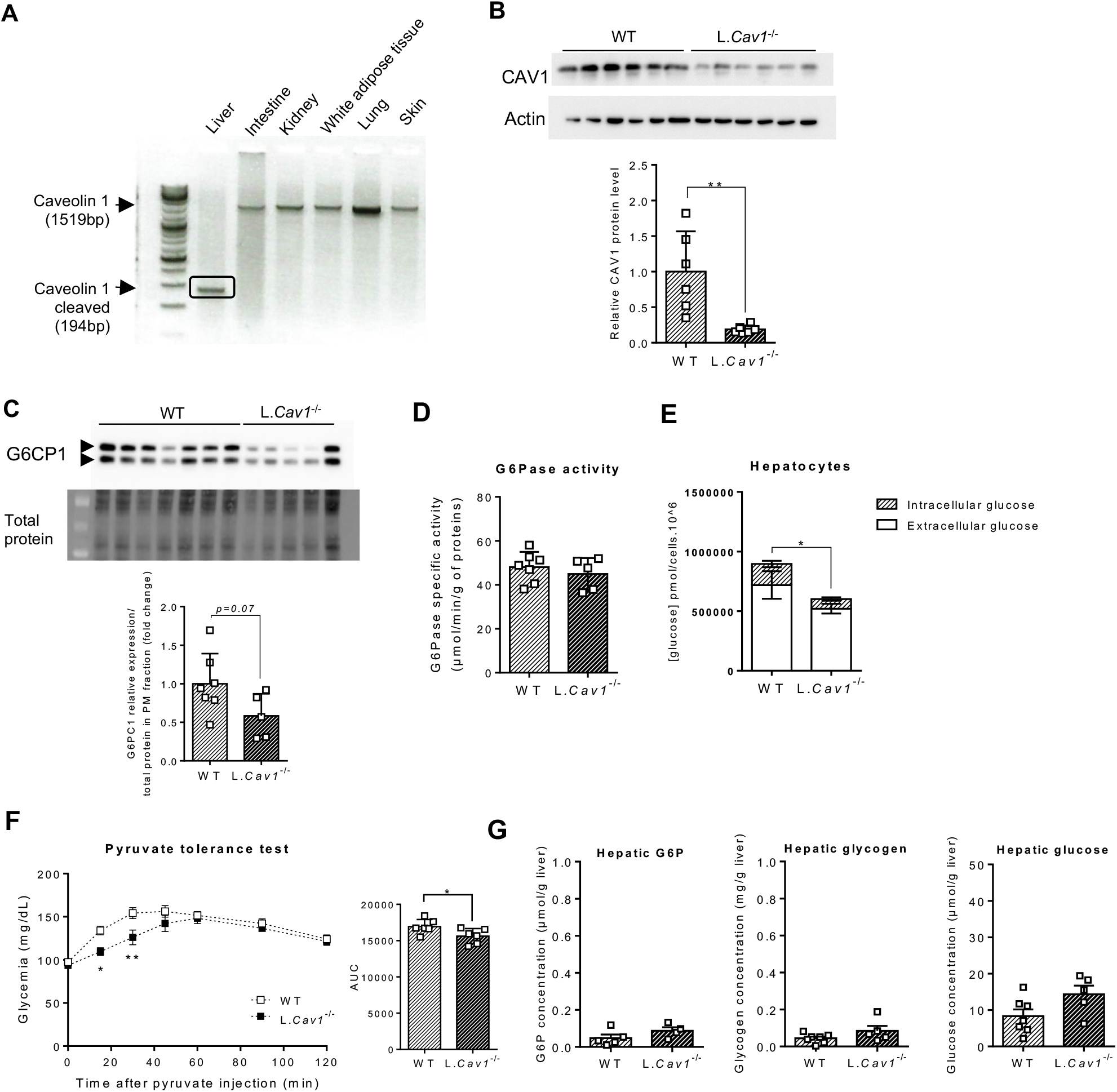
Lack of caveolin-1 in the liver reduces hepatic glucose production. (**A**) Representative image of PCR genotype analysis on the basis of tail DNA in *L*.*Cav1*^*-/-*^ mice after tamoxifen injection in different tissue. (**B**) Representative Western blot analysis and quantification of CAV1 accumulation in *L*.*Cav1*^*-/-*^ and WT mice hepatocyte (n=6/group). (**C**) Representative Western blot analysis and quantification of G6PC1 proteins in liver plasma membrane in *L*.*Cav1*^*-/-*^ (n=5) and WT mice (n=7). (**D**) G6Pase activity in *L*.*Cav1*^*-/-*^ (n=5) and WT mice (n=7) liver. (**E**) Intracellular and extracellular glucose produced from pyruvate and lactate in *L*.*Cav1*^*-/-*^ and WT mice hepatocyte (n=6). Hatched bars represent extracellular glucose while white bars represent intracellular glucose. (**F**) Pyruvate tolerance test. Blood glucose levels were measured for 120 min after a pyruvate intraperitoneal injection (n=6/group). (**G**) Hepatic glucose, G6P and glycogen content in mice liver (n=5-7). Data are expressed as mean ± SEM. Significant differences from WT mice are indicated as *P < 0.05; **P < 0.01. Unpaired Student’s t-test were performed on male 16h-fasted mice (standard diet). For pyruvate tolerance test, two-way ANOVA followed by post-hoc test was used.

### L.Cav1^-/-^ mice are protected against diabetes development under a High Fat High Sucrose diet

We then assessed the physiological consequences of suppressing this vesicular pathway of HGP. After 22 weeks of STD diet, glucose, G6P and glycogen levels were comparable between 6h-fasted L.*Cav1*^-/-^ and WT mice (**Figure 2A**). Moreover, there were no differences in plasma glucose and insulin levels in 6h-fasted mice fed a STD diet (**Figure 2B**). Interestingly, L.*Cav1*^-/-^ mice fed a STD diet maintained their plasma glucose levels after a 16h fast despite a lower hepatic glucose production (**Figure 2C**). To assess a possible compensatory increase of glucose production by the intestine and kidneys, we performed a glutamine tolerance test but could not reveal any increase in extrahepatic glucose production (data not shown). These data show that despite reduced HGP, L.Cav1^-/-^ mice do not increase glycogen storage, suggesting that the lack of hepatic CAV1 doesn’t inhibit post-absorptive glycogenolysis, but only reduces fasting gluconeogenesis.

**Figure 2:**
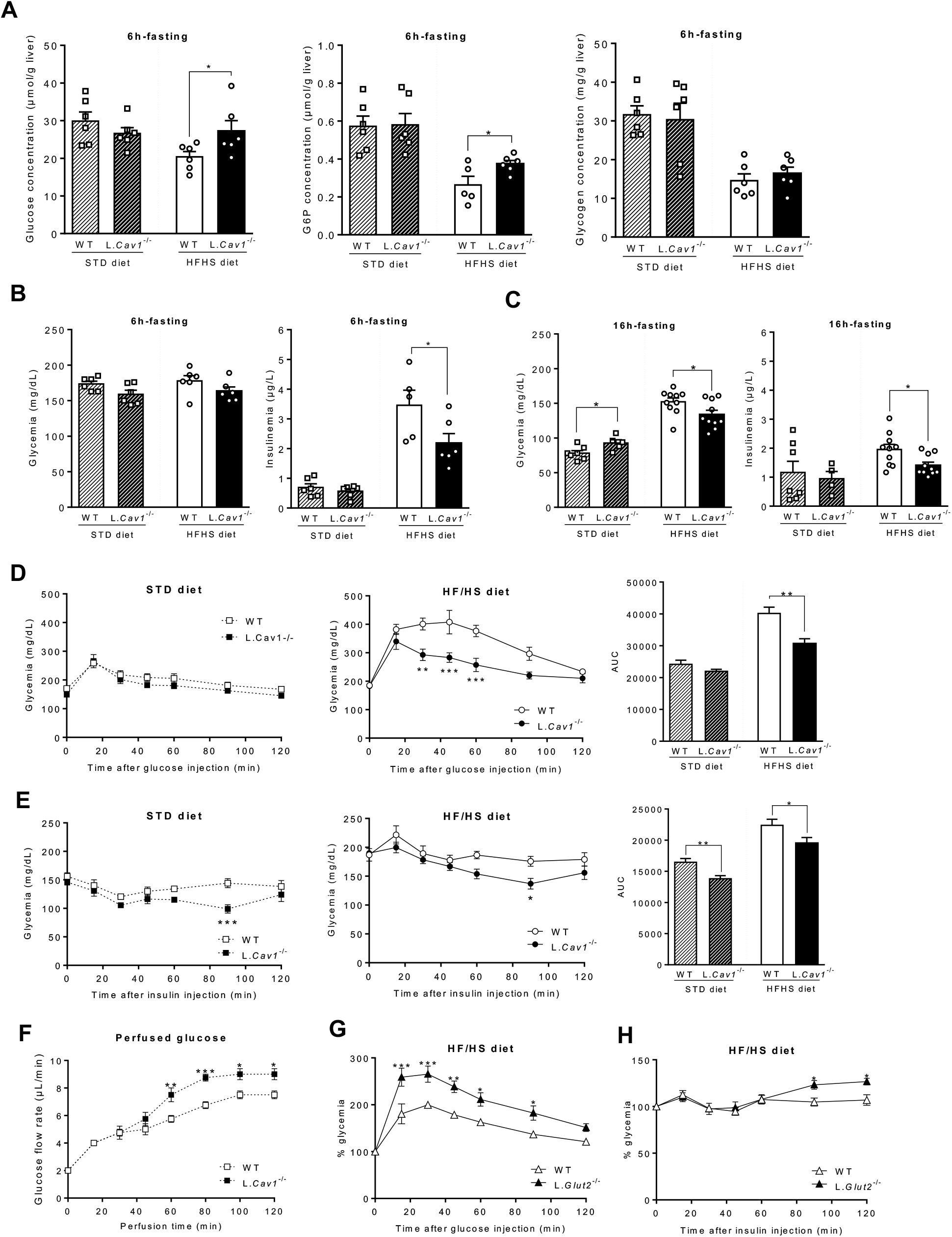
Protection from diabetes in L.*Cav1*^*-/-*^ mice fed a high fat/high sucrose diet. (**A**) Hepatic glucose, G6P and glycogen content in 6h-fasted *L*.*Cav1*^*-/-*^ (white bars) and WT (black bars) mice fed a STD (hatched bars) or HFHS (solid bars) diet (n=6/group). **(B)** Blood glucose and plasma insulin levels in 6h-fasted *L*.*Cav1*^*-/-*^ and WT mice (n=6/group). (**C**) Blood glucose and plasma insulin levels in 16h-fasted *L*.*Cav1*^*-/-*^ and WT mice (n=6/group). Glucose (**D**) and insulin (**E**) tolerance tests (with corresponding AUC) performed in 6h-fasted *L*.*Cav1*^*-/-*^ (black symbols) and WT mice (white symbols) mice fed a STD (squares) or HFHS (circles) diet (n=6/group). **(F)** Glucose infusion under hyperinsulinemia condition, adjusted every 10 min to keep euglycemia in 5h-fasted *L*.*Cav1*^*-/-*^ (white squares) and WT (black squares) mice fed a STD diet (n=4/group). (**G-H**) Glucose (**D**) and insulin (**E**) tolerance tests performed in 6h-fasted *L*.*Glut2*^*-/-*^ (black triangles) and WT (white triangles) mice fed a HFHS diet (n=9-10/group). Data are expressed as mean ± SEM. Two-way ANOVA followed by post-hoc test was used for glucose tolerance test (D), insulin tolerance test (E) and perfused glucose (F). Student’s t-test was performed for glycemia (A/B), insulinemia (A/B) and AUC (D/E). Significant differences from WT mice are indicated as *P < 0.05; **P < 0.01; ***P<0.001.

While glucose tolerance of mice fed a STD diet was not changed by hepatic *Cav1* deletion, a slight improvement of insulin sensitivity was measured in L.*Cav1*^-/-^ mice fed a STD diet compared to WT (**Figure 2D-E**). We performed a hyperinsulinemic euglycemic clamp to confirm this result. The amount of glucose infused to maintain euglycemia during the clamp was higher in L.*Cav1*^-/-^ compared to WT mice demonstrating that hepatic *Cav1* deletion improved glycemic control even in a physiological state (**Figure 2F**).

The first anti-diabetic drug treatment prescribed, Metformin, is thought to exert its primary antidiabetic action through the suppression of HGP^4^. We thus wondered whether the decrease of hepatic gluconeogenesis induced by hepatic *Cav1* invalidation might protect mice from diabetes development. When fed a HFHS diet, 6h-fasted L.*Cav1*^-/-^ mice presented higher hepatic G6P levels compared to WT mice (**Figure 2A**), consistent with a decrease in glucose production at the level of G6Pase. Furthermore, L.*Cav1*^-/-^ mice had lower insulin levels compared to WT (2.2µg/L ± 0.3 vs 3.5µg/L ± 0.5, p=0.05) despite equal plasma glucose levels (**Figure 2B**). These results suggest that liver specific deletion of *Cav1* protects mice against the hyperinsulinemia induced by HFHS diet. Consistently, plasma glucose and insulin levels (1.4µg/L ± 0.11 vs 1.9µg/L ± 0.17, p=0.01) were lower in 16h-fasted L.*Cav1*^-/-^ compared to WT mice fed a HF/HS diet (**Figure 2C**). Glucose and insulin tolerance tests highlighted the improved glucose tolerance and insulin sensitivity of L.*Cav1*^-/-^ mice fed a HF/HS diet compared to WT (**Figure 2D-E**).

Altogether, these results suggest that decreasing fasting HGP by targeting the CAV1-dependent vesicular pathway of glucose production protects against HFHS diet-induced deregulations of glucose metabolism and improves glycemic control even in a physiological state.

Interestingly, L.*Glut2*^-/-^ mice did not produce glucose from glycogenolysis in the post-absorptive state but exhibited a normal HGP during fasting thanks to the CAV1-dependent vesicular pathway of glucose production^12^. We thus investigate whether decreasing HGP in the post-absorptive state (i.e. in L.*Glut2*^-/-^ mice) had the same protective effect against diabetes development than decreasing HGP in the fasting state (in L.*Cav1*^-/-^ mice). L.*Glut2*^-/-^ mice fed a HFHS diet showed impaired glucose tolerance and insulin sensitivity on the contrary to L.*Cav1*^-/-^ mice (Figure 2G-H).

*L*.*Cav1*^*-/-*^ *mice are weakly protected against obesity development under a High Fat High Sucrose diet* We previously showed that blunting HGP protects against weight gain^7^. We then questioned whether L.*Cav1*^-/-^ mice could also be protected against obesity. We measured during 6 months the weight gain of mice fed either STD or HFHS diet and showed that hepatic *Cav1* deletion was associated with a lower weight gain when mice were fed a HFHS diet but not when mice were fed a STD diet, without any difference in food intake (**Figure 3A**). Accordingly, metabolic exploration by magnetic resonance revealed a significant decreased fat mass percentage in L.*Cav1*^-/-^ mice fed a HFHS diet in contrast to WT mice (**Figure 3B**). Concomitantly to the development of obesity, HFHS diet is known to induce hyperlipidemia and hepatic steatosis. No changes in plasma triglyceride and cholesterol levels (**Figure 3C**) or in hepatic triglyceride and cholesterol accumulation (**Figure 3D**) were observed between groups. At the molecular level, hepatic qPCR analyses highlighted increases in gene expression of the master regulators of *de novo* lipogenesis *Srebp1c* (Sterol regulatory element binding protein 1c) and *Chrebp* (Carbohydrate-responsive element binding protein) in the liver of L.*Cav1*^-/-^ compared to WT mice fed a HFHS diet (**Figure 4A**). Furthermore, hepatic *Cav1* deletion induced higher *Acc* (Acetyl-CoA carboxylase), *Scd1* (Stearoyl-CoA desaturase 1) and *Elovl6* (Elongation of very long chain fatty acid protein 6) mRNA levels without changes in *Fas* (Fatty Acid Synthase) gene expression. In parallel, qPCR analyses showed higher expression levels of genes involved in lipid oxidation. We measured higher *Ppara* (Peroxysome proliferator activated receptor alpha), *Acox1* (Peroxisomal acyl-coenzyme A oxidase 1), *Acadl* (Acyl-CoA dehydrogenase) and *Cyp4a10* (Cytochrome p450 4A10) mRNA levels, in the liver of L.*Cav1*^-/-^ compared to WT mice fed a HFHS diet (**Figure 4B**). No differences in the expression of genes controlling de novo lipogenesis and lipid oxydation were measured in the liver of mice fed a STD diet.

**Figure 3:**
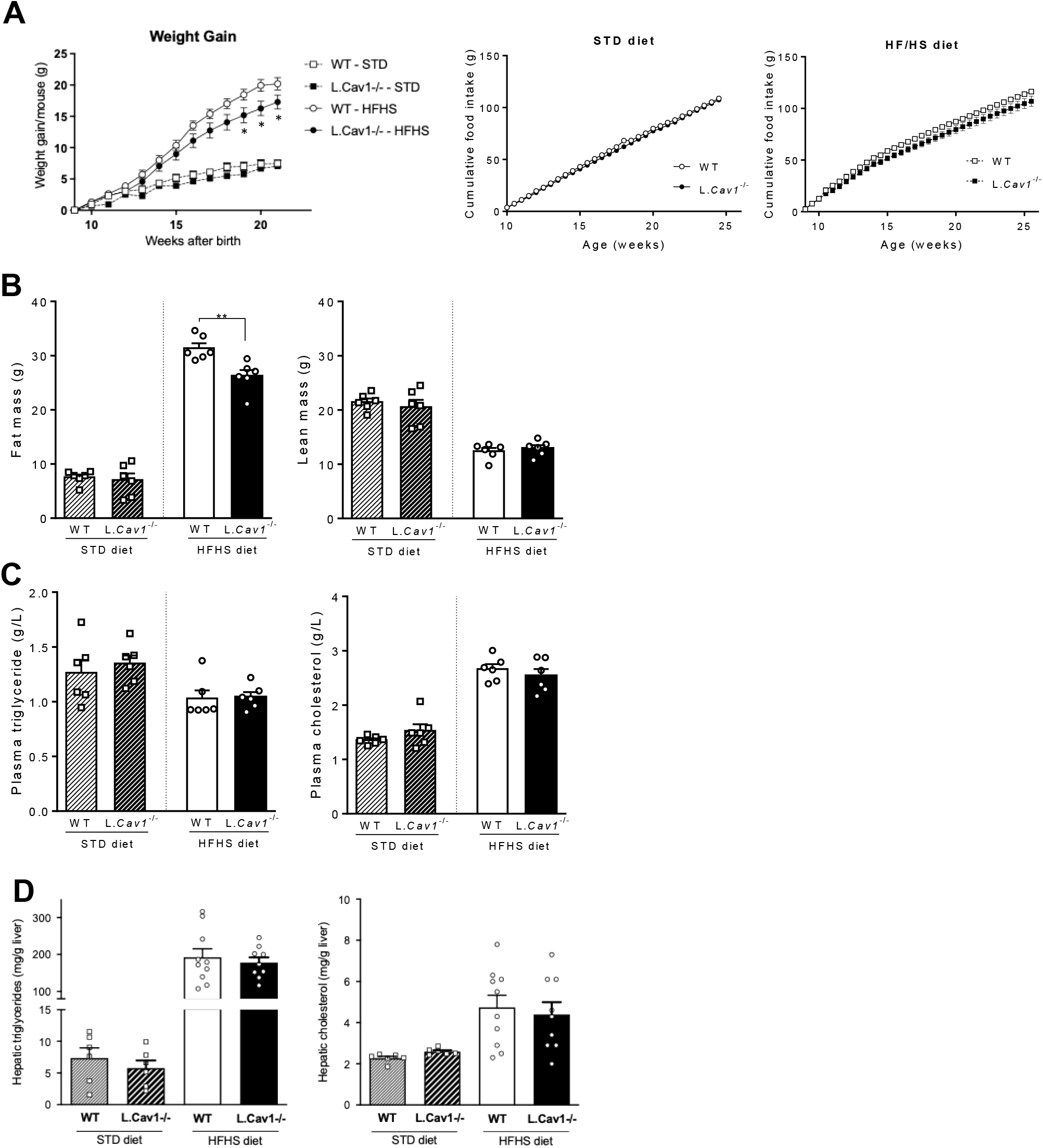
L.*Cav1*^*-/-*^ mice fed high fat/high sucrose diet weakly resist to obesity. (A) Cumulated weight gain (left panel) in *L*.*Cav1*^*-/-*^ (black symbols) and WT (white symbols) mice fed a STD (square symbols) or HFHS (circle symbols) diet. Cumulative food intake (right panel) (n=6/group) (B) %Fat mass / body weight and %Lean mass / body weight obtained by resonance magnetic spectrometer of *L*.*Cav1*^*-/-*^ (white bars) and WT (black bars) mice fed a STD (hatched bars) or a HF/HS (plains bars) diet (n=6/group). (**C**) Plasmatic triglyceride and cholesterol levels in 6h-fasted mice (n=6/group). (**D**) Hepatic triglyceride and cholesterol contents in 6h-fasted mice (n=6/group). Data are expressed as mean ± SEM. Two-way ANOVA followed post-hoc test was used for weight gain, food intake (A). Student’s t-test was performed for fat/lean mass (B), plasmatic triglyceride and cholesterol concentration (C), hepatic triglyceride and cholesterol content (D). Significant differences from WT mice are indicated as *P < 0.05; **P < 0.01; ***P<0.001.

**Figure 4:**
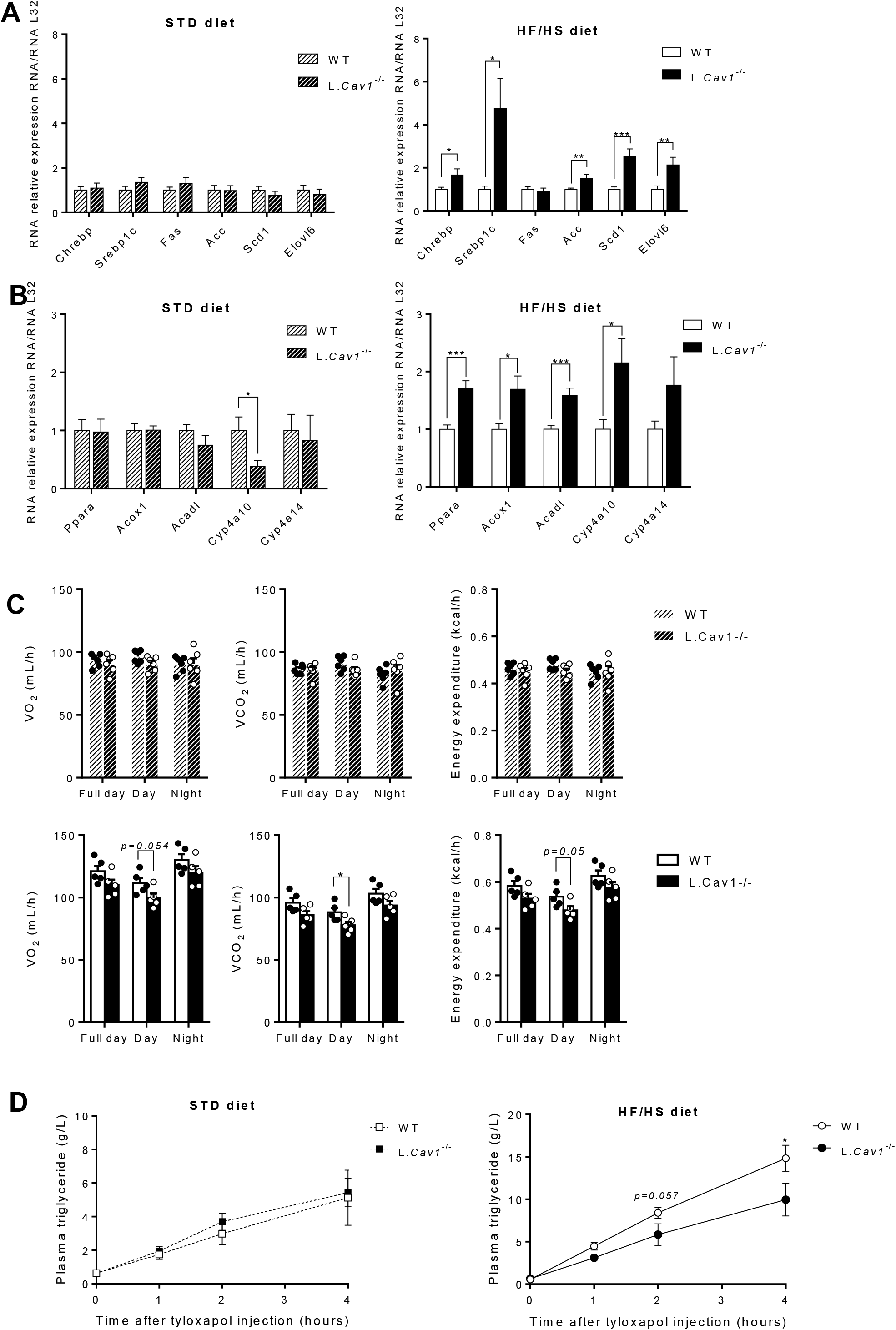
Lack of caveolin-1 in the liver affects hepatic gene expression and intestinal lipid absorption. Relative mRNA levels of genes involved in *de novo* lipogenesis (**A**) and lipid oxidation (**B**) in the liver of 6h-fasted mice (n=6/group STD diet; n=10/group HFHS diet). (**C**) Whole body oxygen consumption, carbon dioxide production and energy expenditure in mice fed a STD diet (upper panel) or a HFHS diet (lower panel) (n=6/group). (**D**) Plasma triglyceride levels measured at 0, 1, 2, 3 and 4h after lipoprotein lipase injection and oil gavage in 6h-fasted mice fed a STD diet (left panel; n=5) or a HFHS diet (right panel; n=8). Data are expressed as mean ± SEM. Two-way ANOVA followed post-hoc test was used for intestinal lipid absorption (D). Student’s t-test was performed for mRNA levels (A/B) and VO_2_, VCO_2_, energy expenditure (C). ANCOVA (Analysis of Covariance) adjusted for lean mass was used on VO_2_, VCO_2_ and energy expenditure (data not shown). Significant differences from WT mice are indicated as *P < 0.05; **P < 0.01; ***P<0.001.

To address whether changes in energy expenditure contributed to the lower weight gain of L.*Cav1*^-/-^, we performed indirect calorimetry measurements. Our data showed decreased VO_2_, VCO_2_ and energy expenditure during the day in L.*Cav1*^-/-^ mice fed a HFHS diet (**Figure 4C**). Analysis of covariance adjusted for lean mass revealed no effect of genotype on VO_2_, VCO_2_ and energy expenditure. We thus studied whether the protection against weight gain of L.*Cav1*^-/-^ mice was due to an impairment in intestinal lipid absorption. After the injection of a lipoprotein lipase (LPL) inhibitor followed by oil gavage, we measured plasma triglyceride levels along time. Plasma triglyceride levels were lower in L.*Cav1*^-/-^ mice fed a HFHS diet (1.5-fold induction in WT mice 4h after injection of LPL inhibitor compared to L.*Cav1*^-/-^ mice, p<0.05) meaning a worse intestinal lipid absorption in mice lacking hepatic CAV1 (**Figure 4D**). All together, these findings suggest that hepatic CAV1 deficiency limits intestinal lipid absorption, which may provide protection against obesity.

### Decreased hepatic glucose production corrects partially diabetes but not obesity and hepatic steatosis in previously prediabetic and overweight mice

We then investigated whether the decreased HGP induced by hepatic CAV1 deficiency could improve glucose homeostasis and energy metabolism in previously prediabetic and overweight mice. To this purpose, we fed mice with a HFHS diet for 12 weeks prior to induce *Cav1* deletion. The HFHS diet was pursued during 2 additional months before we evaluated metabolic parameters. First, we didn’t observe differences in blood glucose levels, nevertheless we measured lower plasma insulin levels in 6h-fasted L.*Cav1*^-/-OB^ mice in contrast to WT (**Figure 5A**). Accordingly, glucose tolerance and insulin sensitivity were improved in L.*Cav1*^-/-OB^ compared to WT mice (**Figure 5B-C**). No change in hepatic glucose, G6P and glycogen levels were measured in 6h-fasted L.*Cav1*^-/-OB^ compared to WT mice (**Figure 5D**). Interestingly, deleting hepatic CAV1 in previously prediabetic and overweight mice did not induce changes in body weight gain (**Figure 5E**). Moreover, comparable plasma triglyceride and cholesterol levels were measured in L.*Cav1*^-/-OB^ and WT mice (**Figure 5F**). In addition, they showed similar hepatic triglyceride and cholesterol accumulation (**Figure 5G**). Thus, the reduced hepatic glucose production caused by the liver-specific deletion of Cav1 partially corrects the deregulation of carbohydrate metabolism independent of a reduction in body weight.

**Figure 5:**
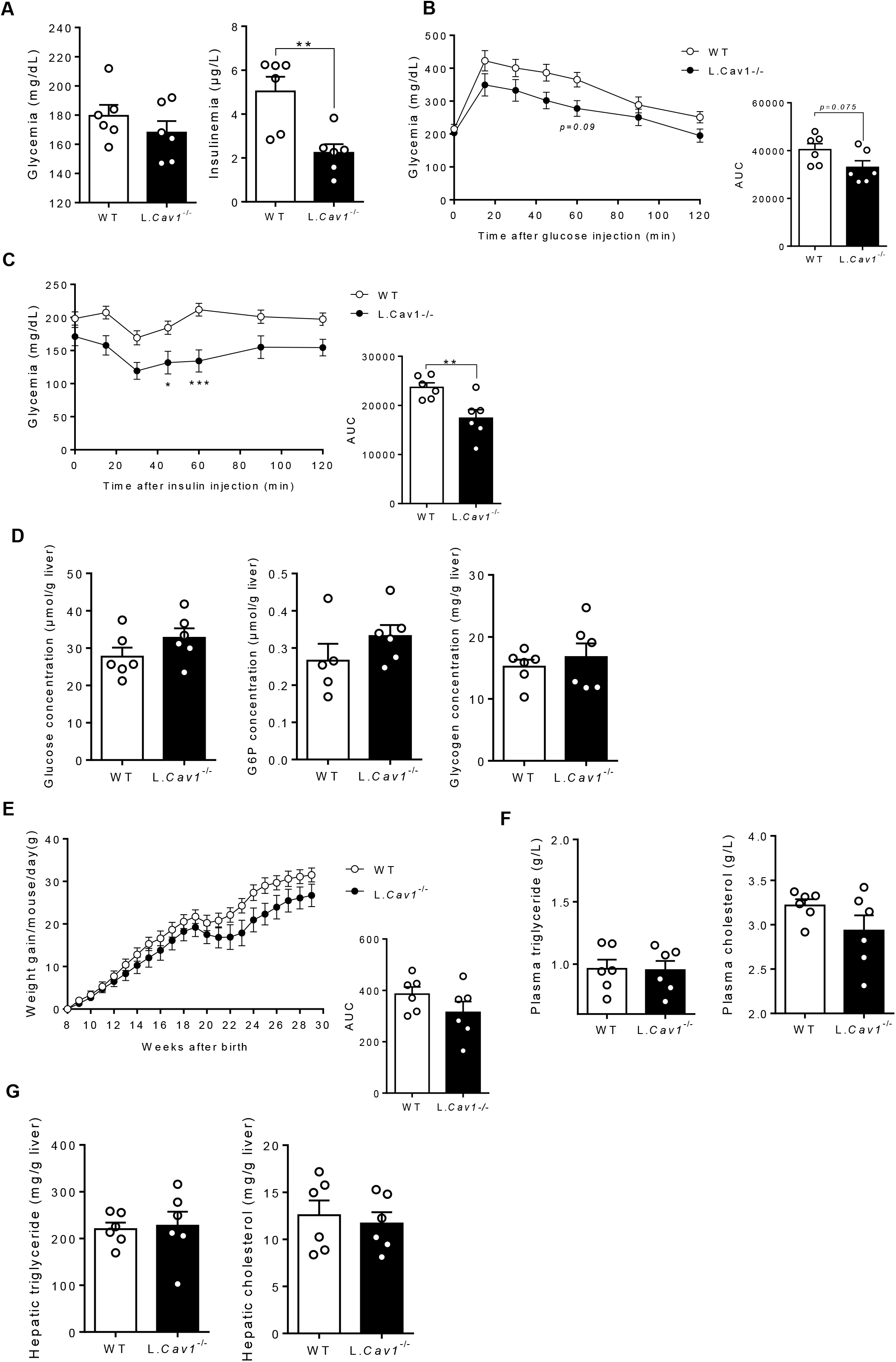
Improvements of glucose balance in previously overweight and prediabetic L.*Cav1*^-/-^ mice. **(A)** Plasma glucose and insulin levels of 6h-fasted *L*.*Cav1*^*-/-*^ (white bars) and WT (black bars) mice (n=6). **(B**) Glucose tolerance test with corresponding AUC performed in 6h-fasted *L*.*Cav1*^*-/-*^ (white symbols) and WT (black symbols) mice (n=6). (**C**) Insulin tolerance test with corresponding AUC performed in 6h-fasted *L*.*Cav1*^*-/-*^ and WT mice (n=6). (**D**) Hepatic glucose, G6P and glycogen contents in 6h-fasted mice. (**E**) Cumulated weight gain with corresponding AUC of *L*.*Cav1*^*-/-*^ and WT mice (n=6). (**F**) Plasma triglyceride and cholesterol levels of 6h-fasted mice (n=6). (**G**) Hepatic triglyceride and cholesterol contents in 6h-fasted mice (n=6). Two-way ANOVA followed by post-hoc test was used for glucose (B) and insulin tolerance (C) tests and cumulated weight gain (E). Student’s t-test was performed for plasma glucose and insulin levels (A), AUC (B, C, E), hepatic glucose, G6P, glycogen contents (D), plasma and hepatic triglyceride and cholesterol levels (F) (n=6/group). Significant differences from WT mice are indicated as *P < 0.05; **P < 0.01; ***P<0.001.

## DISCUSSION

According to current dogma, excessive HGP is a critical event in the development of type 2 diabetes and may play a key role in the onset of fasting hyperglycaemia^1,24^. Recently, we have reported the existence of a post-translational mechanism that regulates G6PC1 localisation and glucose production in hepatocytes and is dependent on CAV1 vesicle trafficking ^8^. By specifically targeting this pathway in the liver, we have here demonstrated its critical role in the regulation of glucose homeostasis.

We first demonstrated that the absence of CAV1 specifically in the liver led to a decrease in HGP *in vitro* and *in vivo* independently of a decrease in G6Pase activity. These results confirm that the decrease in HGP measured in total Cav1^-/-^ is due to the loss of hepatic Cav1 and not the consequence of indirect metabolic pathways (e.g. due to altered adipocyte metabolism in Cav1^-/-^ mice). The remaining hepatocyte glucose production is likely redirected to the GLUT2-dependent pathway, although CAV1 deficiency was not compensated by GLUT2 overexpression in the liver (data not shown). Indeed, deletion of GLUT2 in the liver of L.Cav1^-/-^ mice further reduced glucose production by hepatocytes. The lack of hepatic CAV1 improved insulin sensitivity in the physiological state and protected mice against hyperinsulinemia, glucose intolerance and insulin resistance induced by a HFHS diet. In addition, deleting hepatic *Cav1* in prediabetic and overweight mice corrected hyperinsulinemia, glucose intolerance and insulin resistance. It is noteworthy that the improvements in glucose homeostasis induced by hepatic *Cav1* deletion took place in mice that were still overweight, before any difference in body weight was observed. Interestingly, decreasing HGP in the post-absorptive state, such as in L.*Glut2*^-/-^ mice did not induce the same protective effect. On the contrary to L.*Cav1*^-/-^, L.*Glut2*^-/-^ mice fed a HFHS diet have altered glucose tolerance. Altogether, our results show that decreasing HGP specifically in the fasting state by targeting CAV1 improves glucose homeostasis independently of weight loss.

Mice totally deficient for CAV1 showed an increase in plasma fatty acids and low adiposity due to an inability of the adipose tissue to store lipids^25^. Therefore, *Cav1*^-/-^ mice were resistant to weight gain induced by a high-fat diet. Consistent with previous results, we showed that the lack of hepatic *Cav1* had no effect on body weight gain in mice fed a STD diet^26^. However, we showed that the lack of hepatic *Cav1* slightly protected mice against the weight gain induced by a HFHS diet. Consistent with *in vitro* studies performed in human and mouse hepatocyte cell lines^27^, we observed a higher expression of genes involved in *de novo* lipogenesis (including the master regulators *Srebp1c* and *Chrebp*) in L.*Cav1*^-/-^ mice fed a HFHS diet compared to WT mice. In parallel to the increase in lipogenesis, we measured an increase in the expression of genes involved in lipid oxidation (including its master regulator PPAR*α*), which may participate to the decrease in hepatic lipid content. The induction of these two opposite pathways could explain why no difference in the amount of hepatic lipid contents was observed between the two genotypes.

L.*Cav1*^-/-^ mice were protected against the weight gain induced by HFHS diet. Our data suggest that this protection might partly be due to a decrease intestinal lipid absorption. Previous study suggested that hepatic CAV1 deficiency blocked bile acid export from mouse hepatocytes, thereby limiting their release into the intestine^28^. This decreased release of bile acids in the intestine could explain the lipid malabsorption measured in L.*Cav1*^-/-^ mice. In addition, deleting CAV1 in the liver of previously obese and diabetic mice did not induce weight loss despite an improvement of glucose homeostasis. We thus suggest that CAV1 might independently control lipid homeostasis and HGP.

In conclusion, our results provide evidence that the absence of the caveolin-1-dependent vesicular pathway, which is involved in the post-translational mechanism regulating G6PC1 localization and fasting hepatic gluconeogenesis, prevents and corrects the deregulations of carbohydrate metabolism induced by an HFHS diet, independent of body weight reduction. Our results highlight the vesicular pathway of glucose production as a promising therapeutic target for the prevention of type 2 diabetes.

## Acknowledgments

The authors acknowledge the SFR Lyon Est (CNRS UMS3453 - INSERM US7, UCBL1) facilities and particularly thank the members of the “Animalerie Lyon Est Conventionnelle et SPF” for animal care and the members of the ANIPHY platform (Valerie Oréa and Delphine Guyonnet) for assessing body composition and calorimetry experiments. The authors thank the INRAE (AGS), CNRS (GM) and Inserm (AD) for funding their positions.

## Credit Authorship Contribution Statement

**Lorine Da Costa** : Writing – review & editing, Writing – original draft, Methodology, Investigation, Formal analysis, Data curation, Conceptualization. **Cécile Saint-Béat** : Investigation. **Clara Bron** : Investigation. **Arnaud Lang** : Investigation. **Adeline Duchampt** : Investigation, Formal analysis. **Manon Micoud** : Investigation. **Félicie Evrard** : Investigation. **Gilles Mithieux**: Writing – review & editing. **Amandine Gautier-Stein**: Writing – review & editing, Methodology, Formal analysis, Data curation, Conceptualization, Funding acquisition, Supervision, Project administration.

## Funding

This work was supported by a research grant from the French National Research Agency (Projet-ANR-17-CE14-0026).

## Declaration of competing interest

The authors declare no competing interests.

## Data availability

Further information and requests for resources and reagents should be directed to and will be fulfilled by Amandine Gautier-Stein.

## Notes

### Competing Interest Statement

The authors have declared no competing interest.

